## Supplementary Material for "SuperCellCyto: enabling efficient analysis of large scale cytometry datasets"

### Supplementary Note S1

The manual gating strategy employed by Oetjen et al. for the identification of B cell subsets, reproduced from the original manuscript [[1]](https://www.zotero.org/google-docs/?1g1RcQ):

- Pre-B cell (CD20-): CD45+ CD19+ CD20- CD10+
- Immature B cells (CD20+): CD45+ CD19+ CD20+ CD10+
  - Transitional B cells: CD45+ CD19+ CD20+ CD10+ CD27-
- Mature B cells: CD45+ CD19+ CD20+ CD10-
- Naïve Mature B cells: CD45+ CD19+ CD20+ CD10- CD27- CD21hi
- Exhausted/Tissue-like Memory: CD45+ CD19+ CD20+ CD10- CD27- CD21lo
- Memory B cells: CD45+ CD19+ CD20+ CD10- CD27+
- Activated Mature: CD45+ CD19+ CD20+ CD10- CD27+ CD21lo
- Resting Memory: CD45+ CD19+ CD20+ CD10- CD27+ CD21hi
- CD20- CD10-: CD45+ CD19+ CD20- CD10-
- Plasmablast: CD45+ CD19+ CD20- CD10- CD27+ CD38+
- Plasma Cell: CD45+ CD19+ CD20- CD10- CD27+ CD38+ CD138+

### Supplementary Tables

| Dataset | No. of markers | No. of cells | No. of samples | Tissue | Species | Reference |
| --- | --- | --- | --- | --- | --- | --- |
| Levine_32dim | 32 | 265,627 | 2 | Bone Marrow | Human | [[2]](https://www.zotero.org/google-docs/?80mOKP) |
| Samusik_all | 39 | 841,644 | 10 | Bone Marrow | Mouse | [[3]](https://www.zotero.org/google-docs/?V0GEcZ) |
| Oetjen_bcells | 13 | 8,314,260 | 22 | Bone Marrow | Human | [[1]](https://www.zotero.org/google-docs/?tFpiFg) |
| Trussart_cytofruv | 31 | 8,589,739 | 24 (12 independent) | PBMC | Human | [[4]](https://www.zotero.org/google-docs/?CWHKPq) |
| BCR_XL | 24 | 172,791 | 16 (8 independent) | PBMC | Human | [[5]](https://www.zotero.org/google-docs/?vwmDUI) |
| Anti_PD1 | 24 | 85,715 | 20 | PBMC | Human | [[6]](https://www.zotero.org/google-docs/?dFSh5n) |
| CITEseq | 97 | 49,057 | 10 | Bone Marrow | Human | [[7]](https://www.zotero.org/google-docs/?wEwJM8) |

Supplementary Material Table S1. Overview of datasets used to verify the effectiveness of supercells. All datasets have been previously published, and references to these publications can be found in the final column.

| Dataset | Number of cells | Gamma | Number of supercells |
| --- | --- | --- | --- |
| Levine_32dim | 265,627 | 5 | 53,125 |
|  |  | 10 | 26,563 |
|  |  | 15 | 17,709 |
|  |  | 20 | 13,282 |
|  |  | 25 | 10,625 |
|  |  | 30 | 8,854 |
|  |  | 35 | 7,589 |
|  |  | 40 | 6,641 |
|  |  | 45 | 5,903 |
|  |  | 50 | 5,313 |
| Samusik_all | 841,644 | 5 | 168,328 |
|  |  | 10 | 84,164 |
|  |  | 15 | 56,109 |
|  |  | 20 | 42,082 |
|  |  | 25 | 33,666 |
|  |  | 30 | 28,055 |
|  |  | 35 | 24,048 |
|  |  | 40 | 21,042 |
|  |  | 45 | 18,702 |
|  |  | 50 | 16,832 |

Supplementary Material Table S2. The number of supercells generated by various gamma values for Levine_32dim [[2]](https://www.zotero.org/google-docs/?eiK1ee) and Samusik_all [[3]](https://www.zotero.org/google-docs/?yByFM3) datasets.

**A**

| **Number of metaclusters** | **Grid size** |
| --- | --- |
| 15 | 10 |
| 20 | 10 |
| 25 | 10 |
| 30 | 10 |
| 15 | 11 |
| 20 | 11 |
| 25 | 11 |
| 30 | 11 |
| 15 | 12 |
| 20 | 12 |
| 25 | 12 |
| 30 | 12 |
| 15 | 13 |
| 20 | 13 |
| 25 | 13 |
| 30 | 13 |
| 15 | 14 |
| 20 | 14 |
| 25 | 14 |
| 30 | 14 |

**B**

**k**

10

15

20

25

30

Supplementary Material Table S3. Range of parameter values explored for clustering Levine_32dim and Samusik_all datasets using (A) FlowSOM [[8]](https://www.zotero.org/google-docs/?0gGA5R) and (B) Louvain [[9]](https://www.zotero.org/google-docs/?Zuajgq) algorithms. For FlowSOM, a square grid was employed for the Self Organising Map (SOM). The grid size represents the size of the square grid. For example, a grid size of 10 represents a square SOM grid of size 10x10.

**A**

| **Gamma** | **< 0.5** | **0.5 (inc.) - 0.9 (exc.)** | **0.9 (inc.) - 1 (exc.)** | **1** |
| --- | --- | --- | --- | --- |
| 5 | 1 (0.002%) | 0 (0.0%) | 1,546 (3.23%) | 46,376 (96.77%) |
| 10 | 5 (0.02%) | 1,635 (6.31%) | 12 (0.05%) | 24,248 (93.62%) |
| 15 | 7 (0.04%) | 1,328 (7.59%) | 94 (0.54%) | 16,070 (91.83%) |
| 20 | 5 (0.04%) | 1,076 (8.17%) | 152 (1.15%) | 11,941 (90.64%) |
| 25 | 8 (0.08%) | 901 (8.54%) | 178 (1.69%) | 9,462 (89.7%) |
| 30 | 7 (0.08%) | 776 (8.83%) | 196 (2.23%) | 7,811 (88.86%) |
| 35 | 9 (0.12%) | 665 (8.82%) | 202 (2.68%) | 6,663 (88.38%) |
| 40 | 8 (0.12%) | 587 (8.9%) | 207 (3.14%) | 5,797 (87.85%) |
| 45 | 3 (0.05%) | 525 (8.95%) | 197 (3.36%) | 5,140 (87.64%) |
| 50 | 3 (0.06%) | 483 (9.15%) | 193 (3.66%) | 4,599 (87.14%) |

**B**

| **Gamma** | **< 0.5** | **0.5 (inc) - 0.9 (exc.)** | **0.9 (inc.) - 1 (exc.)** | **1** |
| --- | --- | --- | --- | --- |
| 5 | 146 (0.1%) | 11,170 (7.98%) | 51 (0.04%) | 128,599 (91.88%) |
| 10 | 258 (0.35%) | 10,573 (14.28%) | 1,253 (1.69%) | 61,979 (83.68%) |
| 15 | 299 (0.59%) | 8,610 (17.12%) | 2,231 (4.44%) | 39,151 (77.85%) |
| 20 | 305 (0.8%) | 7,211 (18.93%) | 2,579 (6.77%) | 27,999 (73.5%) |
| 25 | 278 (0.91%) | 6,255 (20.42%) | 2,665 (8.7%) | 21,430 (69.97%) |
| 30 | 281 (1.1%) | 5,554 (21.67%) | 2,626 (10.24%) | 17,174 (66.99%) |
| 35 | 268 (1.22%) | 5,027 (22.8%) | 2,628 (11.92%) | 14,127 (64.07%) |
| 40 | 251 (1.3%) | 4,586 (23.71%) | 2,568 (13.28%) | 11,936 (61.71%) |
| 45 | 247 (1.43%) | 4,222 (24.51%) | 2,482 (14.41%) | 10,275 (59.65%) |
| 50 | 239 (1.54%) | 3,898 (25.1%) | 2,417 (15.57%) | 8,973 (57.79%) |

Supplementary Material Table S4. Breakdown of the purity score obtained for the (A) Levine_32dim [[2]](https://www.zotero.org/google-docs/?QAnYE2) dataset and (B) Samusik_all [[3]](https://www.zotero.org/google-docs/?T059cV) dataset across all gamma values. *Inc.* stands for inclusive, while *exc.* represents exclusive.

| **Cytometry Data Cell Type** | **CITEseq Data Cell Type** |
| --- | --- |
| CD16-_NK_cells | CD56brightCD16- NK cells |
| CD16+_NK_cells | CD56dimCD16+ NK cells |
| CD4_T_cells | CD4+ cytotoxic T cells,  CD4+ memory T cells,  Naive CD4+T cells |
| CD8_T_cells | CD8+ central memory T cells,  CD8+ effector memory T cells,  CD8+ naive T cells,  CD8+CD103+ tissue resident memory T cells |
| Mature_B_cells | CD11c+ memory B cells,  Mature naive B cells,  Class switched memory B cells,  Nonswitched memory B cells |
| Monocytes | Classical Monocytes,  Non-classical monocytes |
| pDCs | Plasmacytoid dendritic cells |
| Plasma_B_cells | Plasma cells |
| Pre_B_cells | Small pre-B cell |
| Pro_B_cells | pro-B cells |
| CD34+_HSCs_and_HSPCs | Lymphoid-primed multipotent progenitors,  Megakaryocyte progenitors,  Erythro-myeloid progenitors,  NK cell progenitors,  HSCs & MPPs |

Supplementary Material Table S5. Mapping of cell type labels between Levine_32dim [[2]](https://www.zotero.org/google-docs/?wtx7Qd) cytometry data and the CITEseq data [[7]](https://www.zotero.org/google-docs/?dPSwy9).

| **Cell type** | **Proportion** |
| --- | --- |
| CD4_T_cells | 0.25307149 |
| Monocytes | 0.2025167 |
| CD8_T_cells | 0.19300468 |
| Mature_B_cells | 0.15856561 |
| Pre_B_cells | 0.0588862 |
| CD34+_HSCs_and_HSPCs | 0.04333679 |
| CD16-_NK_cells | 0.03748176 |
| CD16+_NK_cells | 0.02157721 |
| pDCs | 0.01188282 |
| Basophils | 0.01158527 |
| Pro_B_cells | 0.00492398 |
| Plasma_B_cells | 0.00316747 |

Supplementary Material Table S6. Proportions of cell types in Levine_32dim cytometry data [[2]](https://www.zotero.org/google-docs/?THjoCt). Each row corresponds to a specific cell type, with the proportion of cells belonging to that cell.

| **Process** | **Algorithm** | **Resolution** | **Dataset** | **Platform Used** |
| --- | --- | --- | --- | --- |
| Supercells Generation | SuperCellCyto | Single Cells | Anti_PD1,  BCR_XL,  Levine_32dim,  Oetjen_bcells,  Samusik_all,  Trussart_cytofruv | 2022 MacBook Pro (M2 chip, 24GB RAM) |
| Clustering | FlowSOM | Supercells,  Single Cells | Levine_32dim,  Samusik_all | 2022 MacBook Pro (M2 chip, 24GB RAM) |
| Clustering | Louvain | Supercells | Levine_32dim,  Samusik_all | 2022 MacBook Pro (M2 chip, 24GB RAM) |
| Batch Correction | cyCombine | Supercells,  Single Cells | Trussart_cytofruv | 2022 MacBook Pro (M2 chip, 24GB RAM) |
| Clustering | Louvain | Supercells,  Single Cells | Oetjen_bcells | High-Performance Computing (HPC) Platform |
| Clustering | Louvain | Single Cells | Levine_32dim,  Samusik_all | High-Performance Computing (HPC) Platform |
| Batch Correction | CytofRUV | Supercells,  Single Cells | Trussart_cytofruv | High-Performance Computing (HPC) Platform |

Supplementary Material Table S7. The computing platforms used for benchmarking the performance of various analysis processes. Nextflow [[10]](https://www.zotero.org/google-docs/?z5sF7f) pipelines were used to run the processes on the High Performance Computing (HPC) platform. The amount of RAM and CPUs allocated are specified within the Nextflow scripts available on <https://phipsonlab.github.io/SuperCellCyto-analysis/>.

### Supplementary Figures

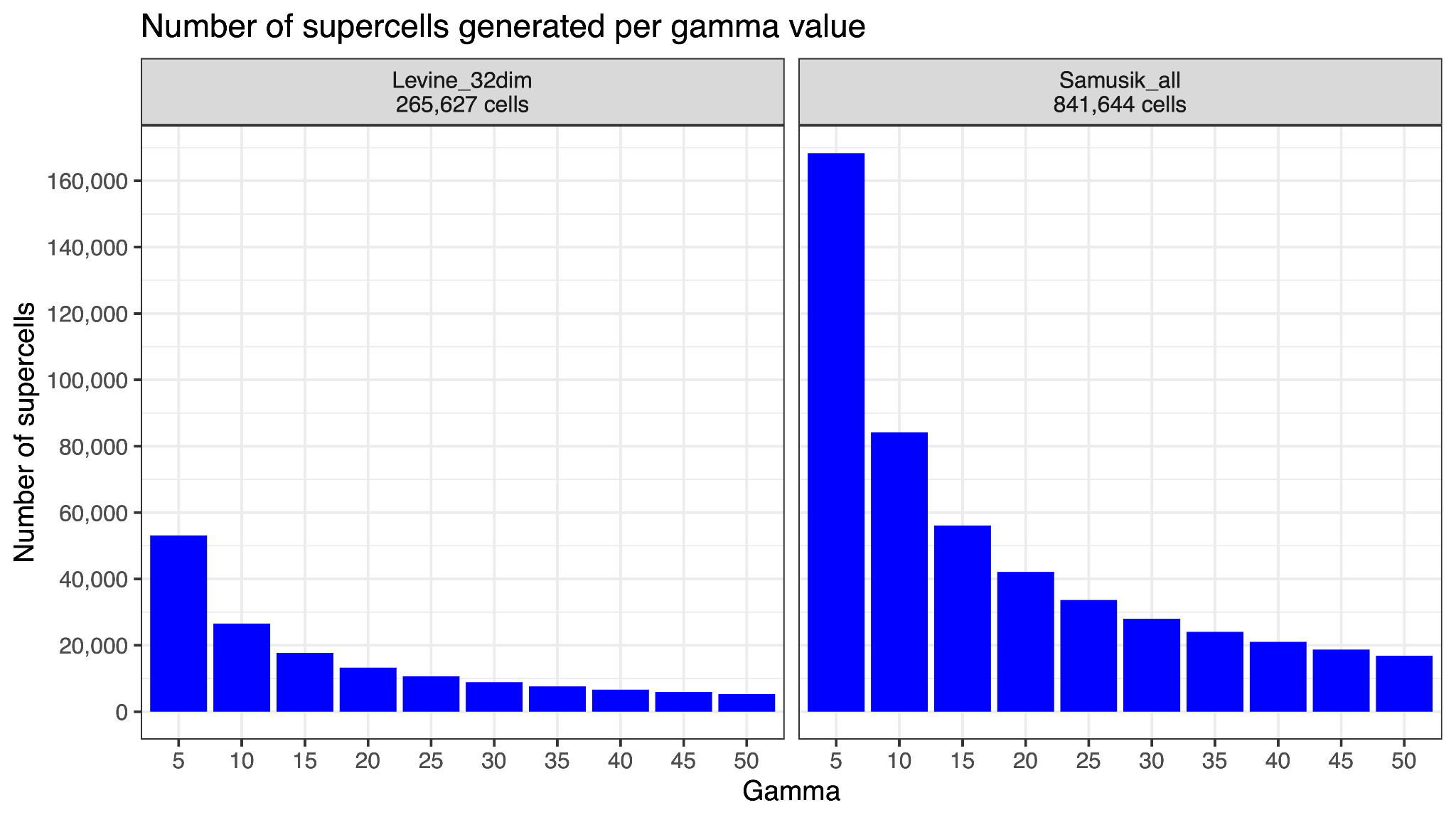

Supplementary Figure S1. Number of supercells generated for Levine_32dim [[2]](https://www.zotero.org/google-docs/?wibLef) and Samusik_all [[3]](https://www.zotero.org/google-docs/?tb1uNJ) datasets across various gamma values.

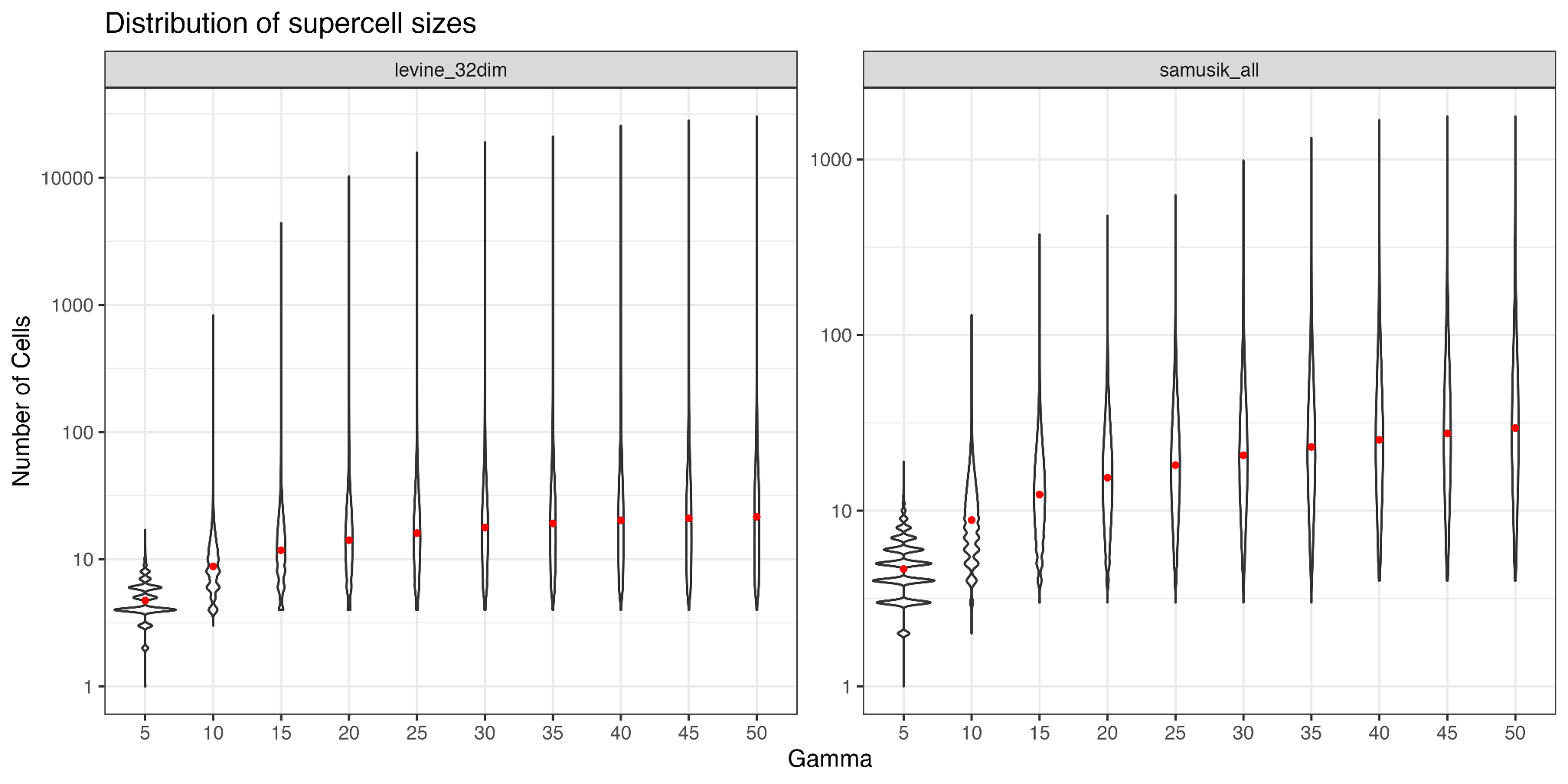

Supplementary Figure S2. Distribution of the number of cells captured in the supercells generated for Levine_32dim [[2]](https://www.zotero.org/google-docs/?q42Vvj) and Samusik_all [[3]](https://www.zotero.org/google-docs/?urEWuZ) datasets using different gamma values. The red dot denotes the mean of the distribution.

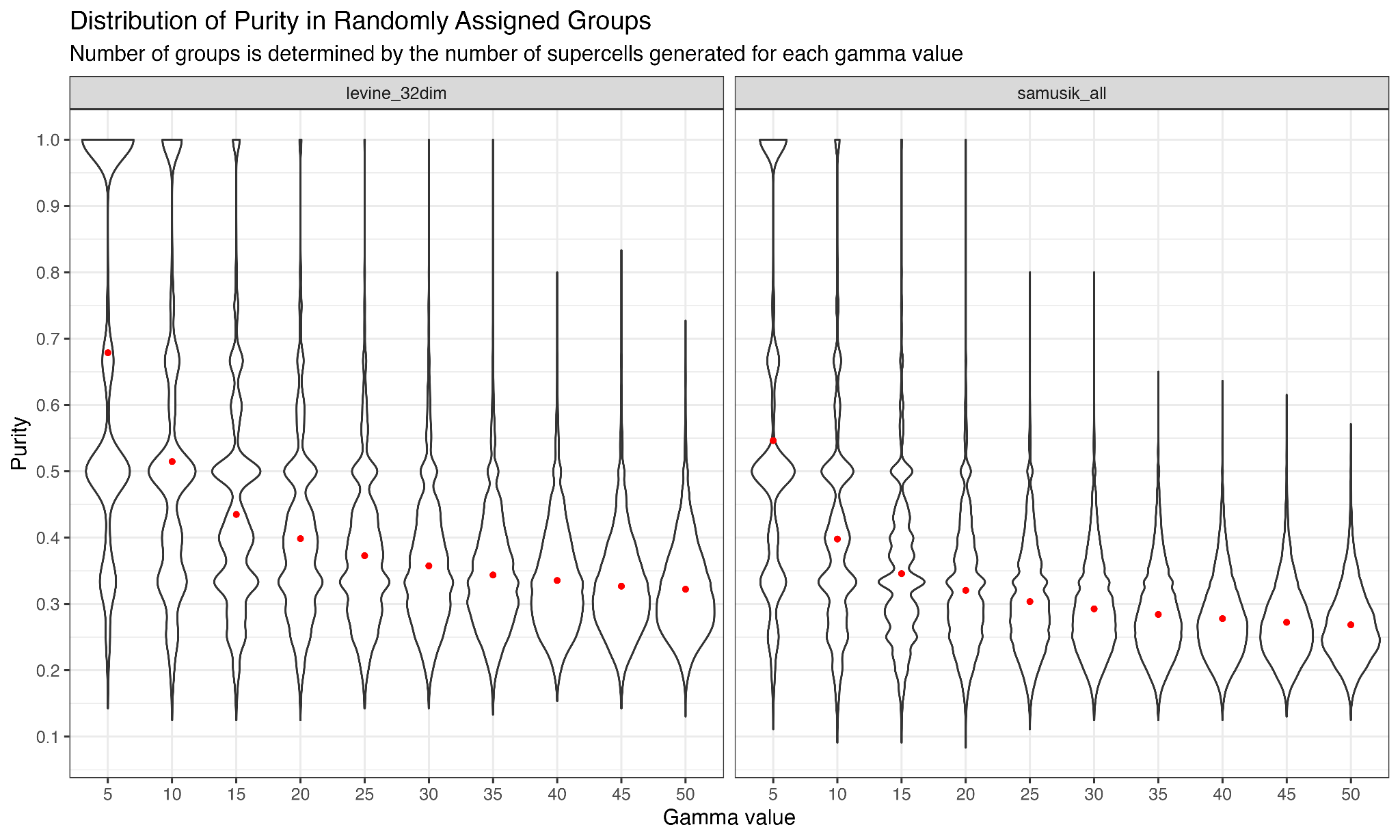

Supplementary Figure S3. Violin plot illustrating the purity score of cell groupings, where cells are randomly assigned into groups. The number of groups corresponds to the number of supercells generated for specific gamma values for a given dataset. See Supplementary Material Figure S1 for the number of supercells generated for a given dataset and gamma value.

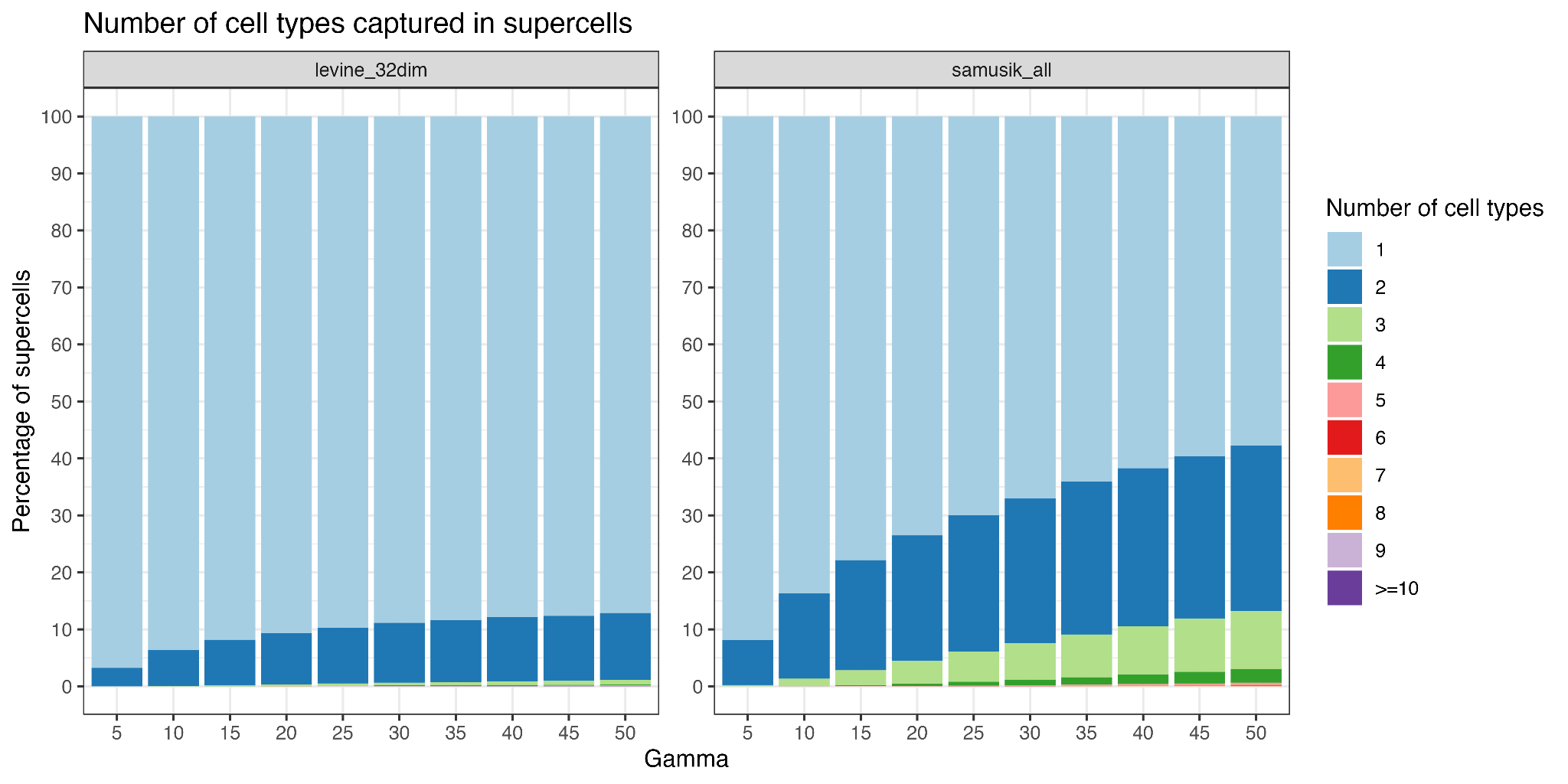

Supplementary Figure S4. Number of different cell types captured in supercells generated for Levine_32dim [[2]](https://www.zotero.org/google-docs/?5vvEJS) and Samusik_all [[3]](https://www.zotero.org/google-docs/?6BUnB4) datasets using different gamma values.

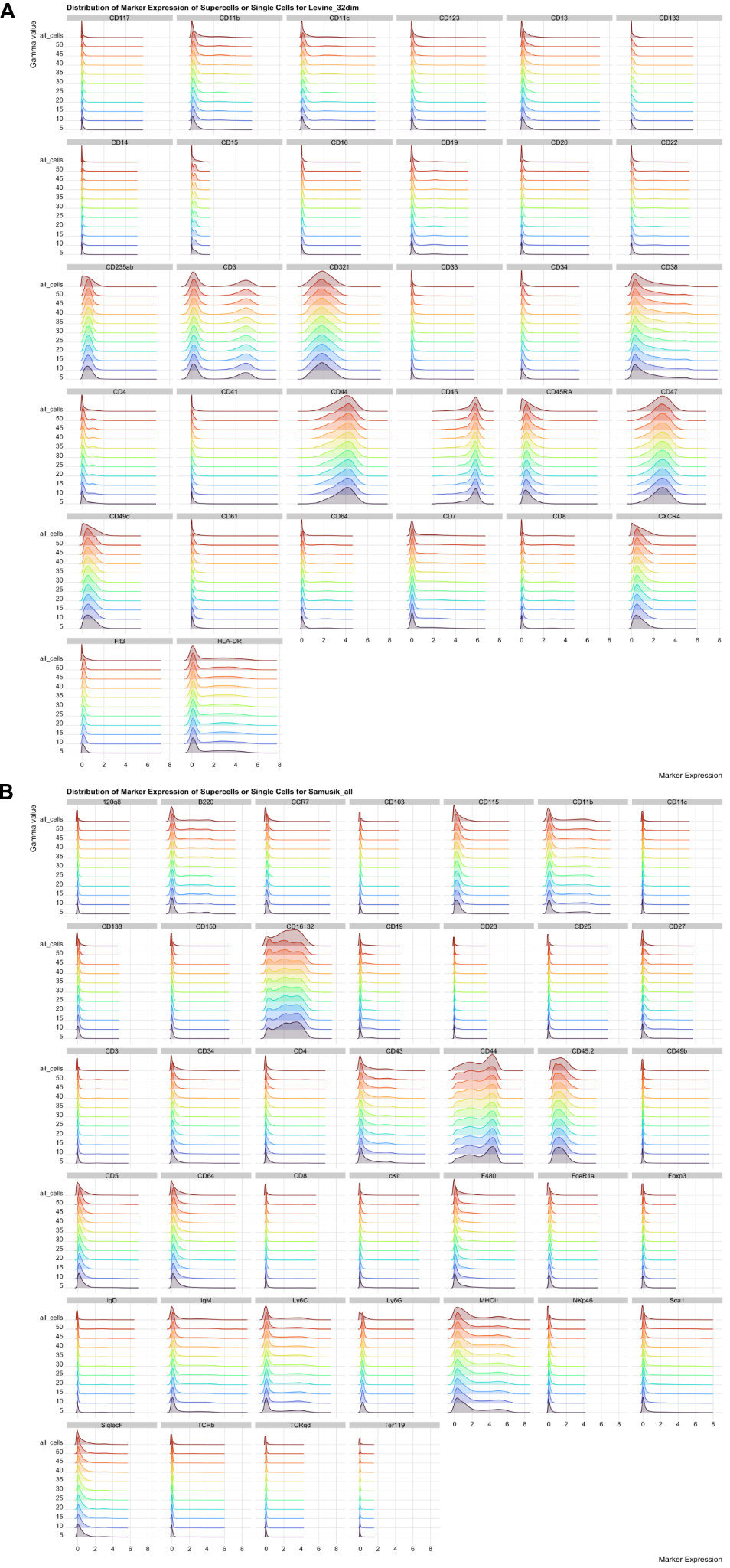

Supplementary Figure S5. Distribution of marker expression for supercells generated using gamma values ranging from 5 to 50 (in increments of 5), and for single cells (denoted by the 'all_cells' row label) for Levine_32dim [[2]](https://www.zotero.org/google-docs/?zl46DV) and Samusik_all [[3]](https://www.zotero.org/google-docs/?OrLJUg) datasets.

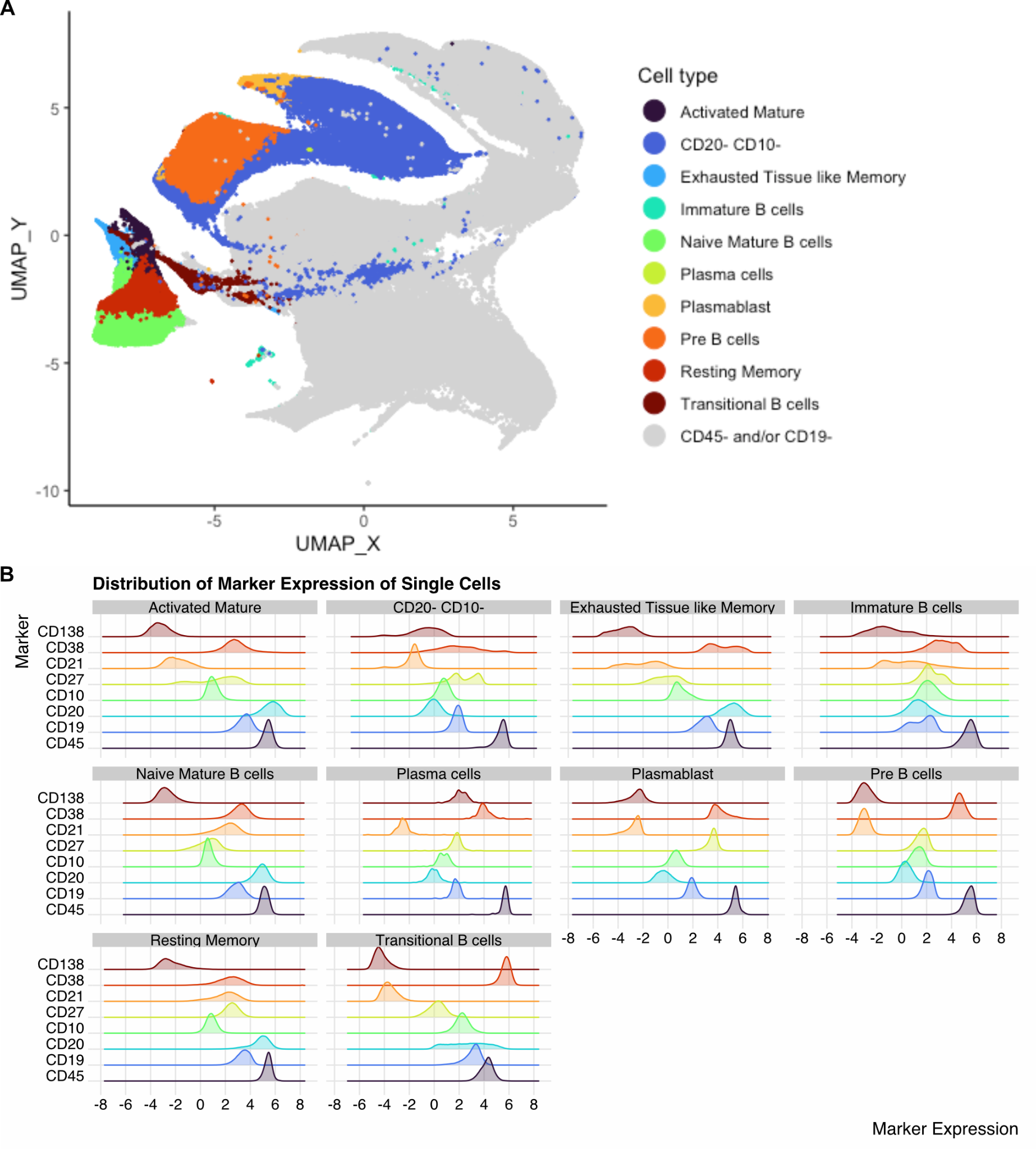

Supplementary Figure S6. UMAP plot and marker expression distribution for Oetjen_bcells data [[1]](https://www.zotero.org/google-docs/?HKwwjz). (A) UMAP plot illustrating supercells generated for the Oetjen_bcells dataset. Each point represents a supercell, and coloured according to the cell type it represents. (B) Distribution of marker expression for single cells in the annotated supercells. Single cells were extracted from annotated supercells, and assigned the corresponding supercell’s annotation.

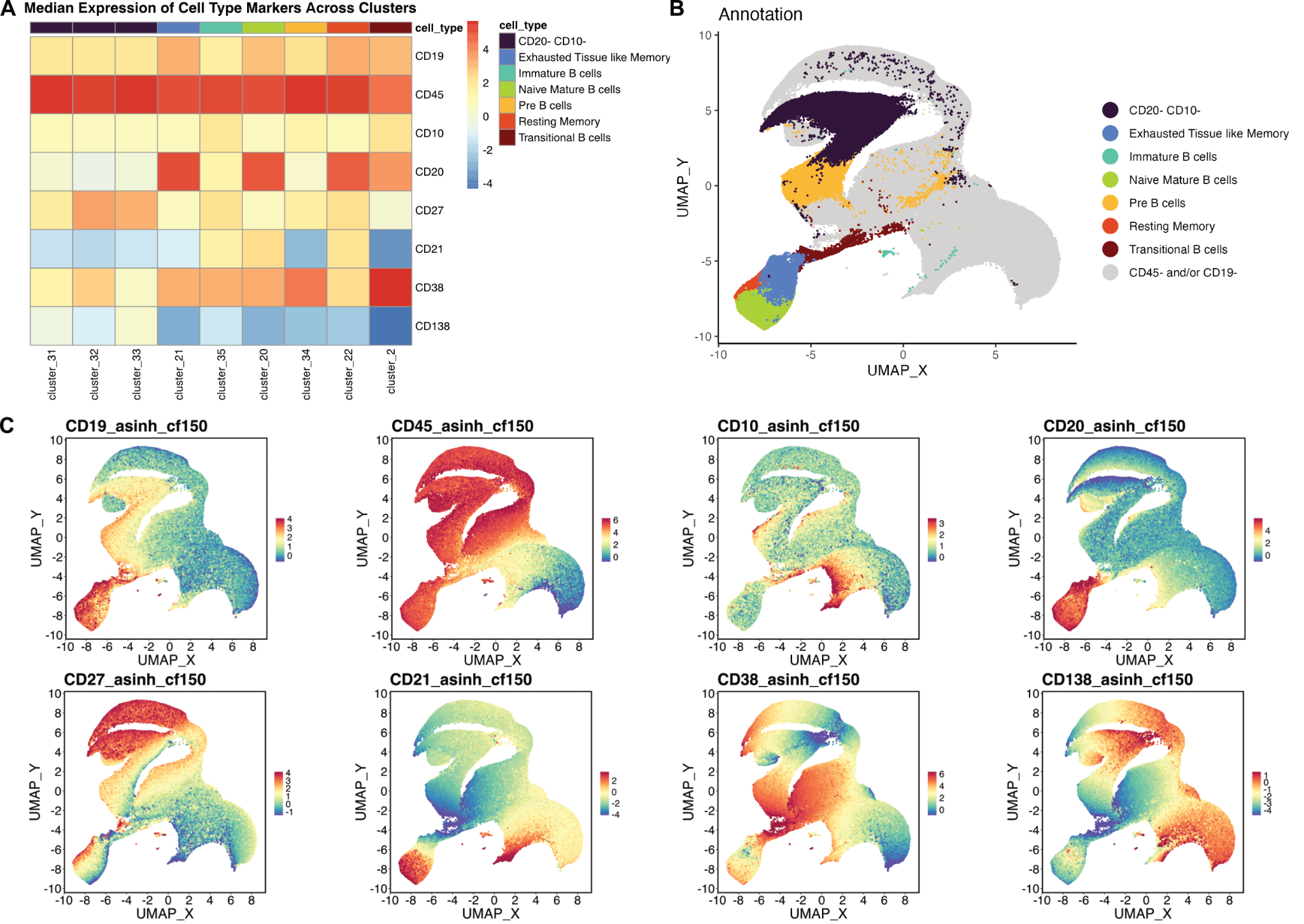

Supplementary Figure S7. Heatmap and UMAP plots for randomly subsampled 415,711 cells from the Oetjen_bcells data [[1]](https://www.zotero.org/google-docs/?NiVbTN). (A) Median expression of cell type markers. Only clusters assigned one of the B cell subsets are shown (B-C) UMAP plots illustrating the B cell subsets identified and the expression of the cell type markers used to annotate the cells.

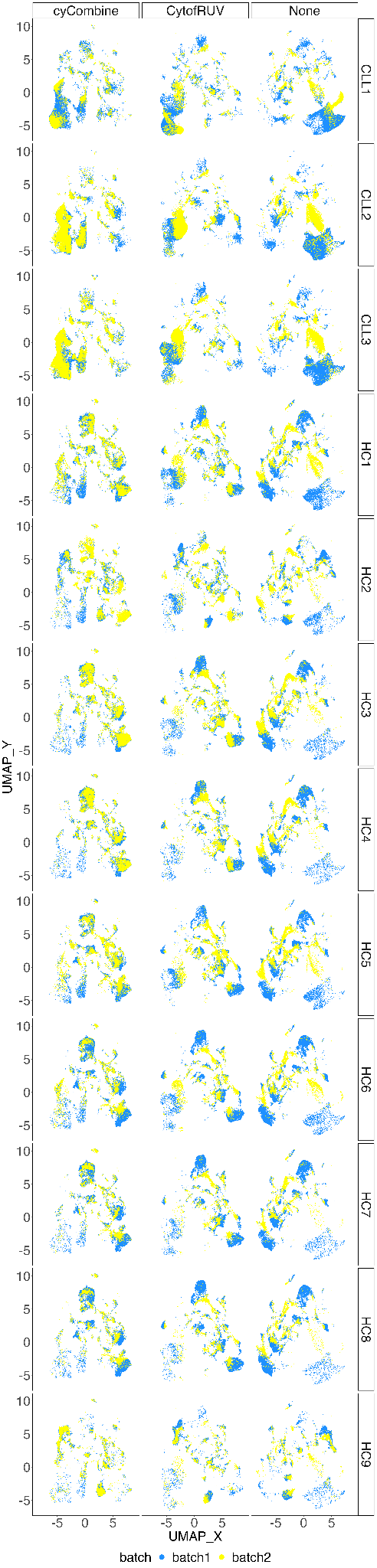

Supplementary Figure S8. UMAP plots of supercells for Trussart_cytofruv dataset [[4]](https://www.zotero.org/google-docs/?Q4EtpG). Each row represents the UMAP plots corresponding to a specific paired sample from a patient, with supercells either corrected using CytofRUV or cyCombine, or left uncorrected.

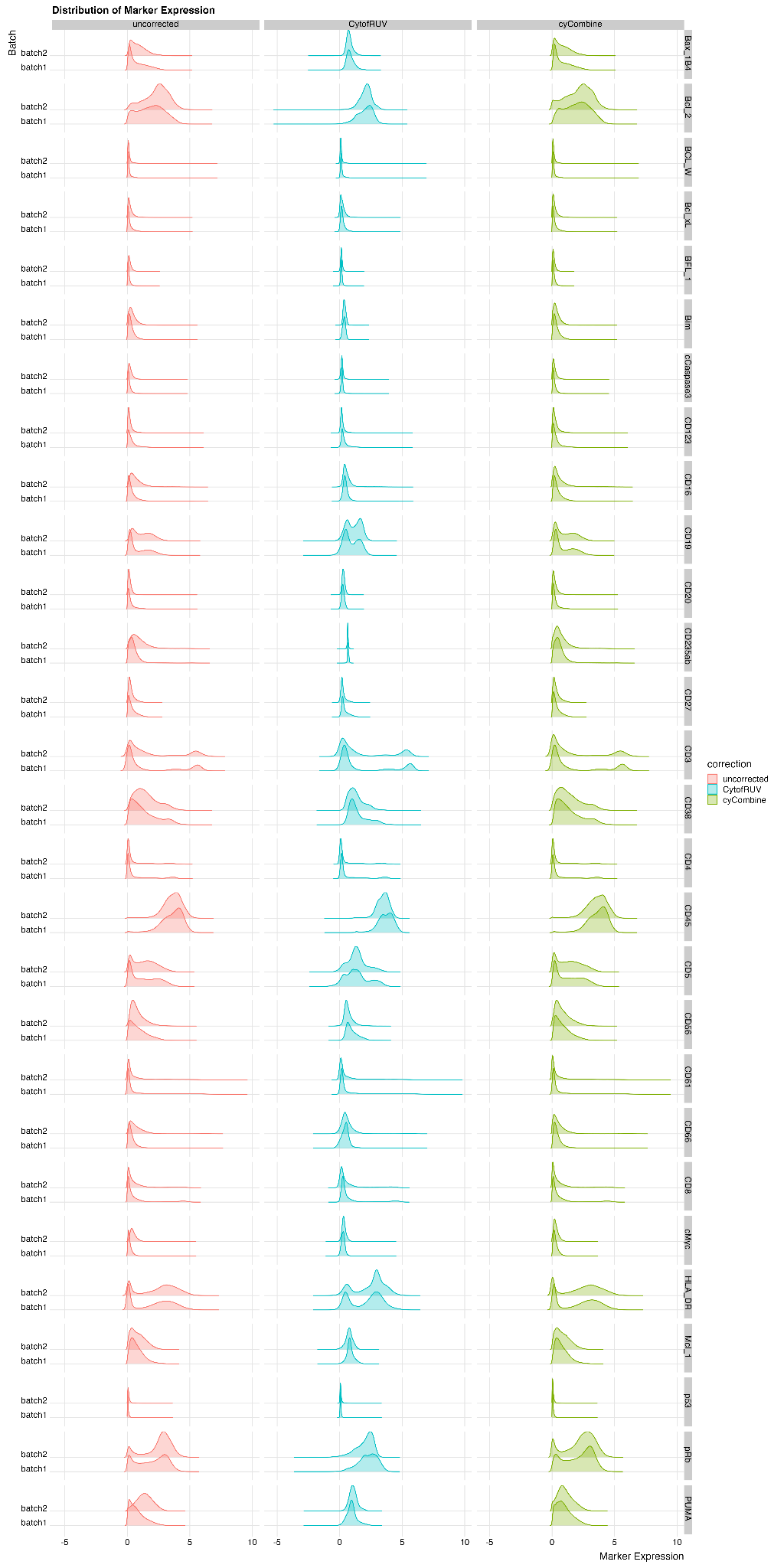

Supplementary Figure S9. Distribution of marker expression for uncorrected, CytofRUV-, and cyCombine-corrected supercells for Trussart_cytofRUV [[4]](https://www.zotero.org/google-docs/?476KMk) dataset.

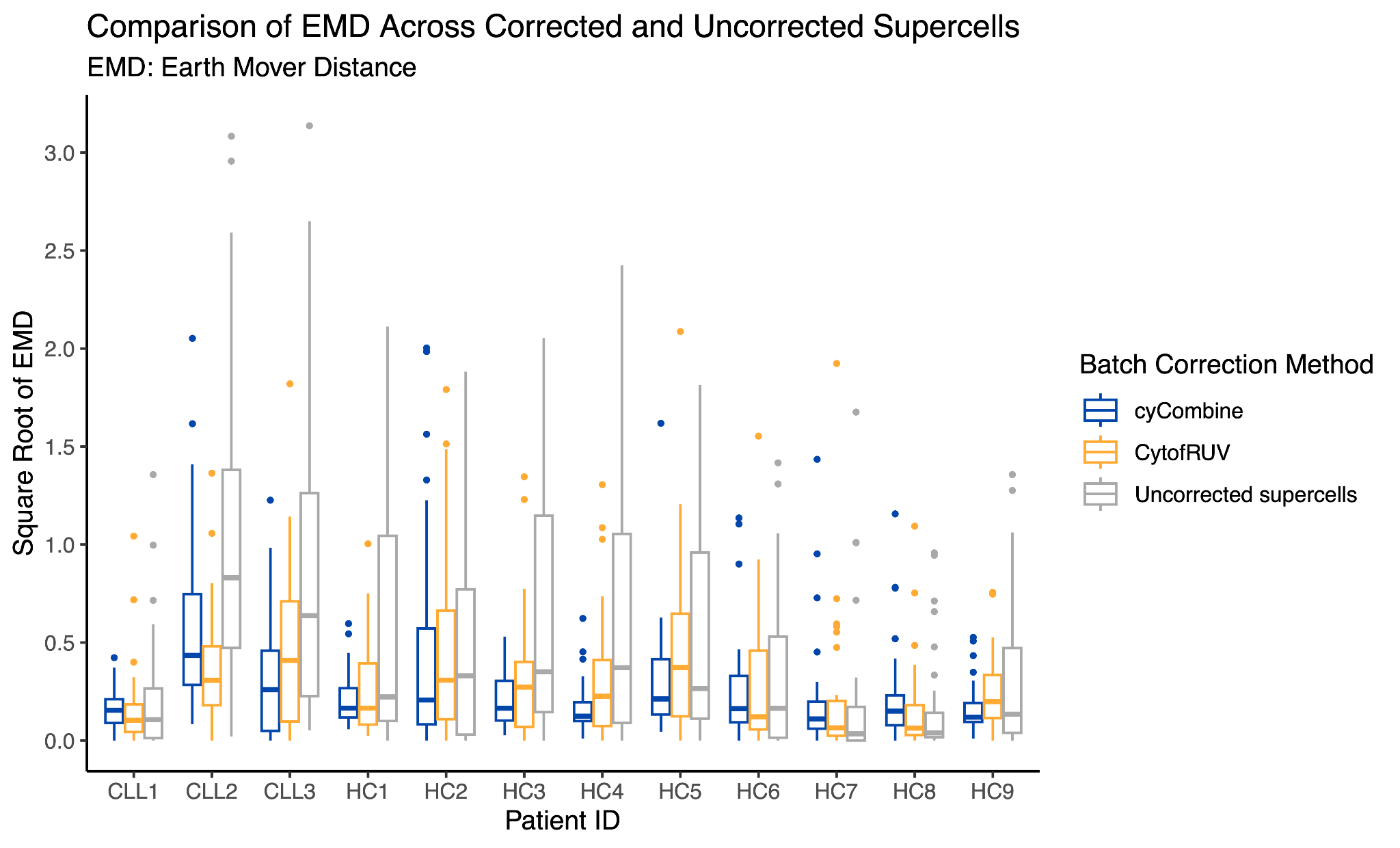

Supplementary Figure S10. A comparison of the Earth Mover Distance (EMD) calculated for all markers before and after the application of batch effect correction for Trussart_cytofRUV dataset [[4]](https://www.zotero.org/google-docs/?VyTZXB). EMD values on the y-axis underwent a square-root transformation to facilitate data visualisation and interpretation.
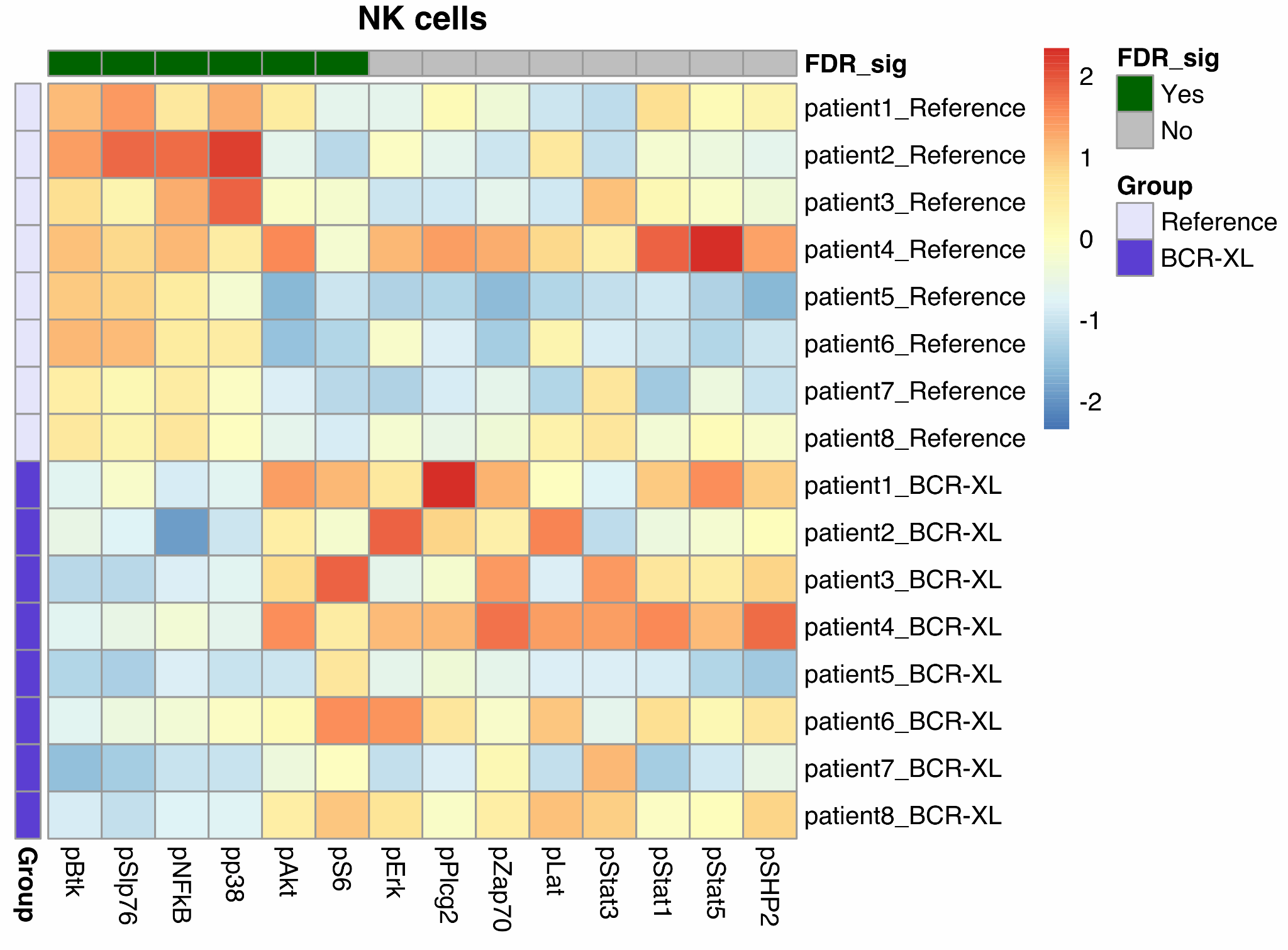

Supplementary Figure S11. Differential expression analysis results generated by Limma [[11]](https://www.zotero.org/google-docs/?u4M8y2) for NK cells in the BCR_XL dataset [[5]](https://www.zotero.org/google-docs/?kwZDBH). The heatmap illustrates the scaled and centred median expression of cell state markers, calculated for each sample across the supercells.

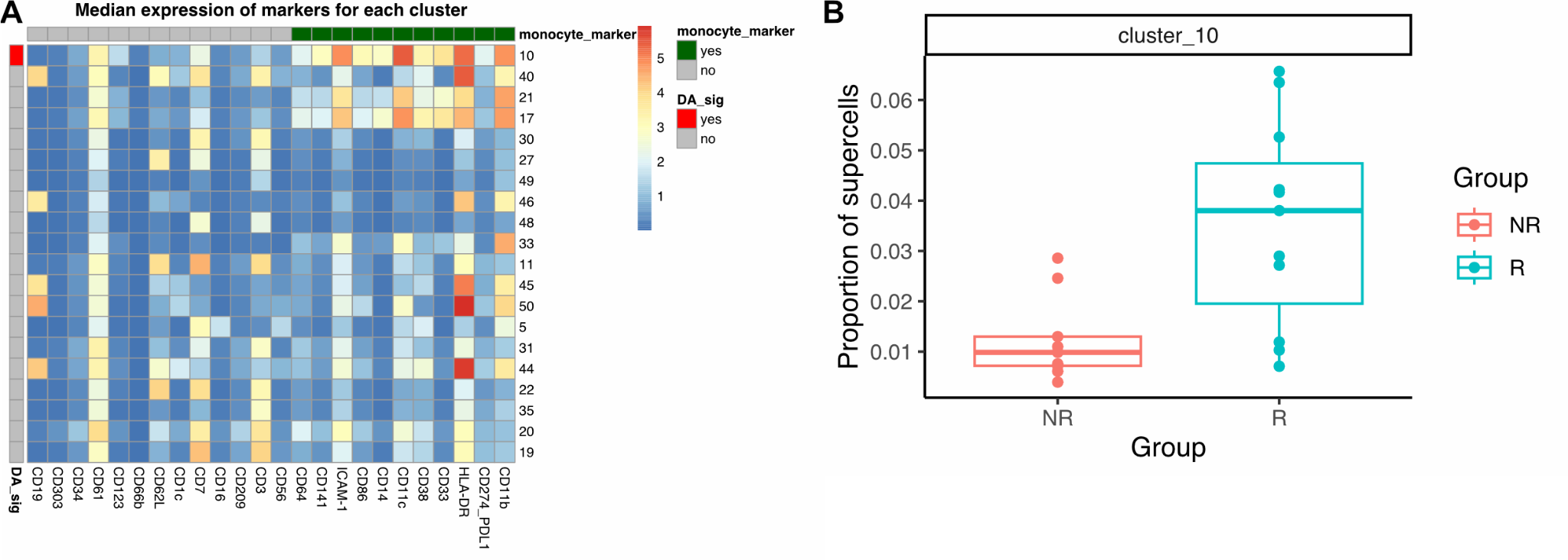

Supplementary Figure S12. Differential abundance analysis for the Anti-PD1 dataset [[6]](https://www.zotero.org/google-docs/?YV5C0k). (A) Heatmap depicting the median expression of markers for each cluster calculated across the supercells. Each marker (column) is annotated based on its function for identifying the rare monocyte subset whose abundance is strongly associated with melanoma patients’ responder status to anti-PD-1 immunotherapy (monocyte_marker). Each cluster (row) is annotated based on the statistical significance of their abundance variation as determined by Propeller (DA_sig, FDR <= 0.05). (B) The proportion of supercells for cluster 10 which represents the rare monocyte subset.

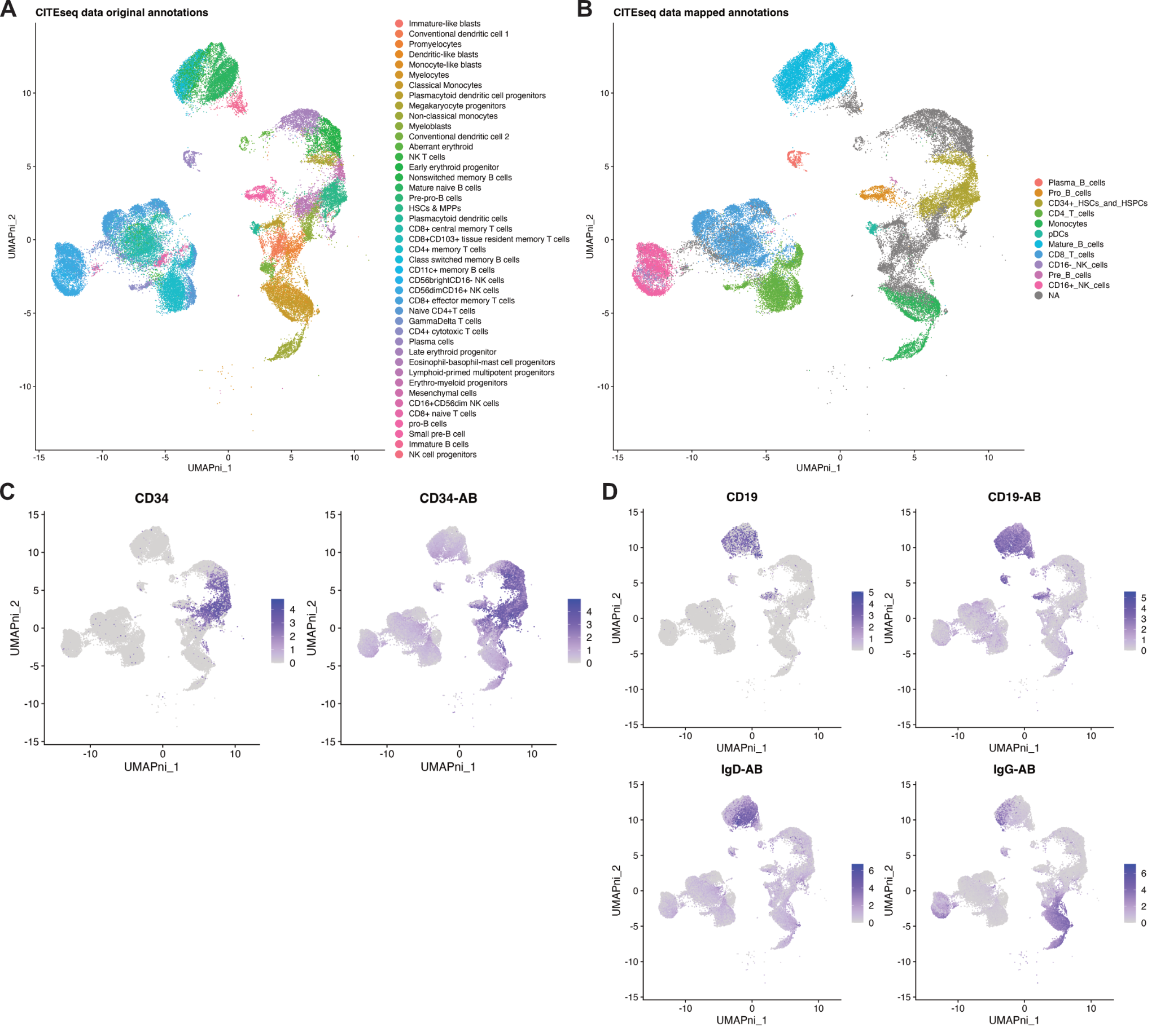

Supplementary Figure S13. UMAP plots of the CITEseq data [[7]](https://www.zotero.org/google-docs/?KhikKS) used in label transfer analysis. Plots are coloured by (A) the original cell type label annotations provided by the authors of the data [[7]](https://www.zotero.org/google-docs/?ugdbgR), (B) the mapped cell type labels for annotating the cytometry data used to calculate accuracy and weighted accuracy metrics, (C) CD34 RNA and antibody (CD34-AB) expressions, (D) CD19 RNA, CD19 (CD19-AB), IgD, and IgG antibodies expressions.

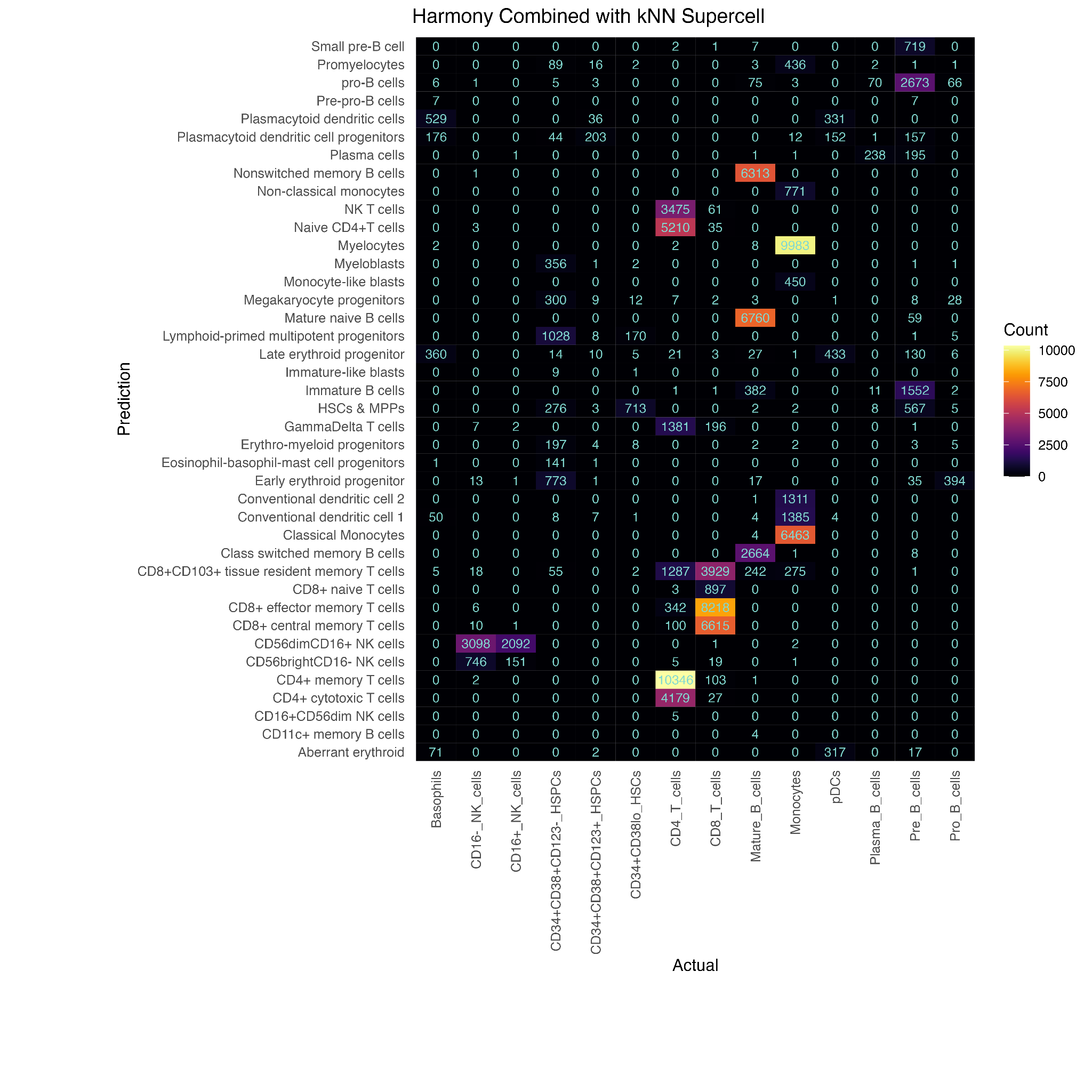
Supplementary Figure S14. Performance evaluation of the cell type label transfer workflow, from a CITEseq dataset to supercells generated for the Levine_32dim cytometry data using Harmony [[12]](https://www.zotero.org/google-docs/?SChm5f) combined with a k-Nearest Neighbour (kNN) classifier.

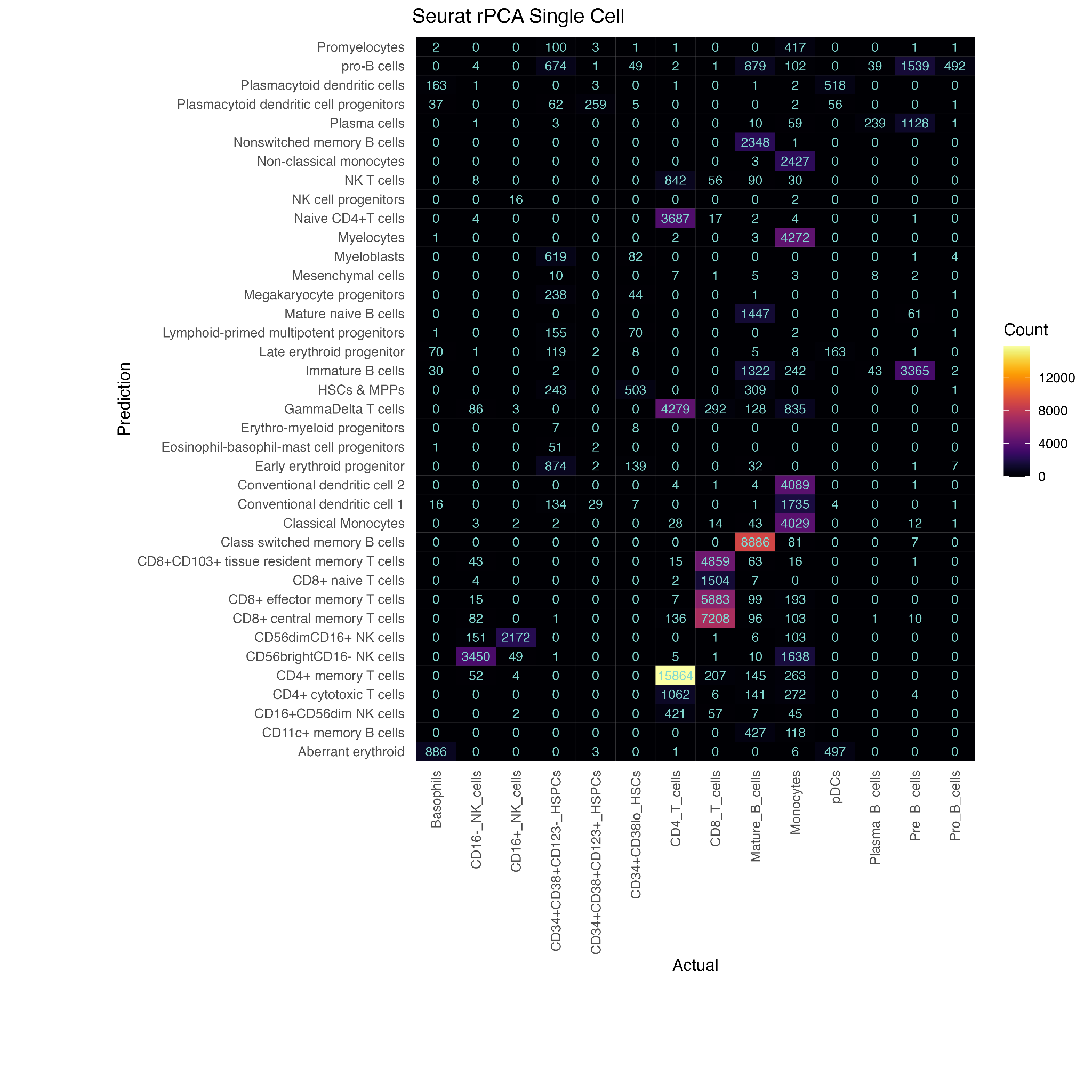
Supplementary Figure S15. Performance evaluation of the cell type label transfer workflow, from a CITEseq dataset to the single cells in the Levine_32dim cytometry data, using Seurat rPCA [[13]](https://www.zotero.org/google-docs/?a3XEPd).

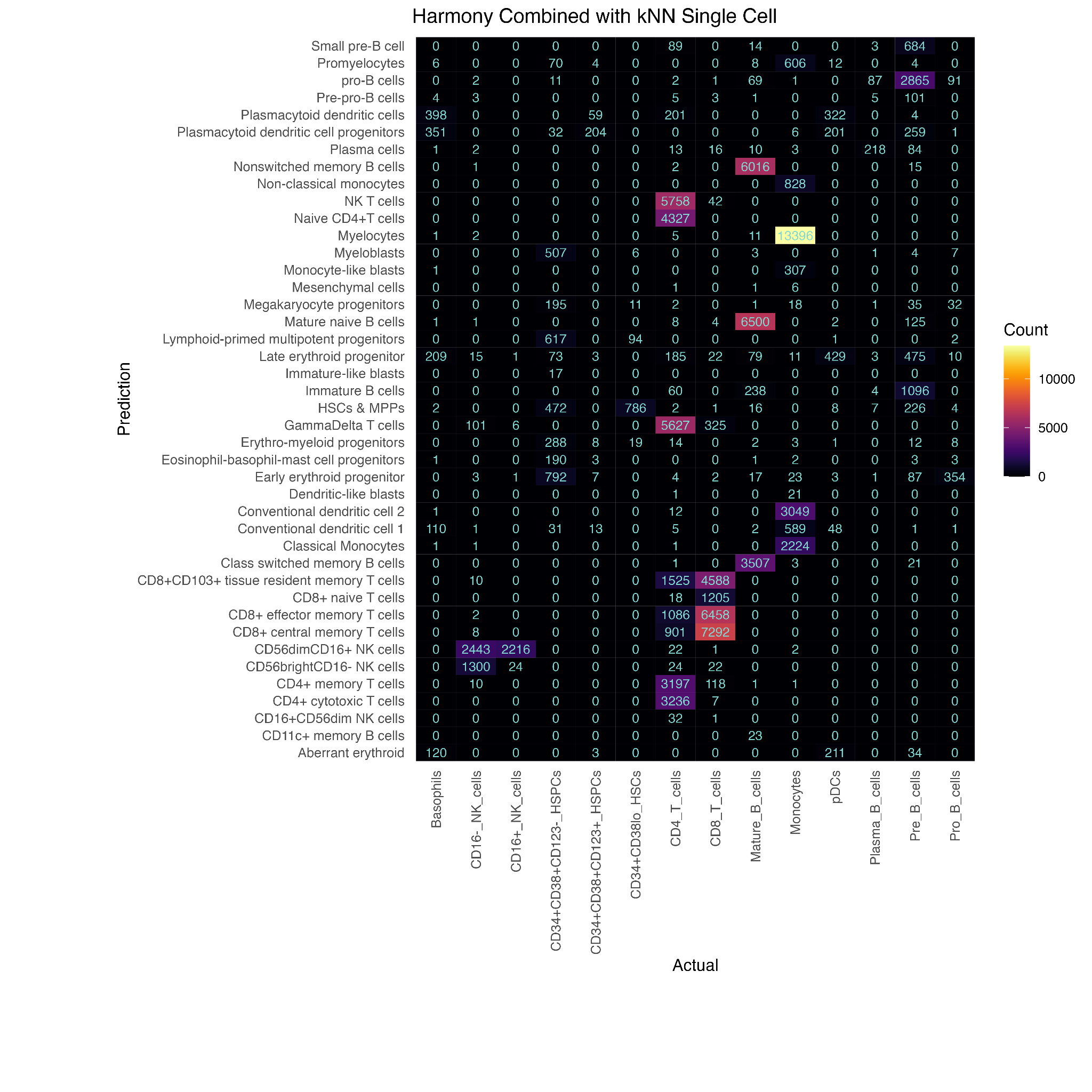
Supplementary Figure S16. Performance evaluation of the cell type label transfer workflow, from a CITEseq dataset to single cells in the Levine_32dim cytometry data using Harmony [[12]](https://www.zotero.org/google-docs/?p2QzbS) combined with a k-Nearest Neighbour (kNN) classifier.

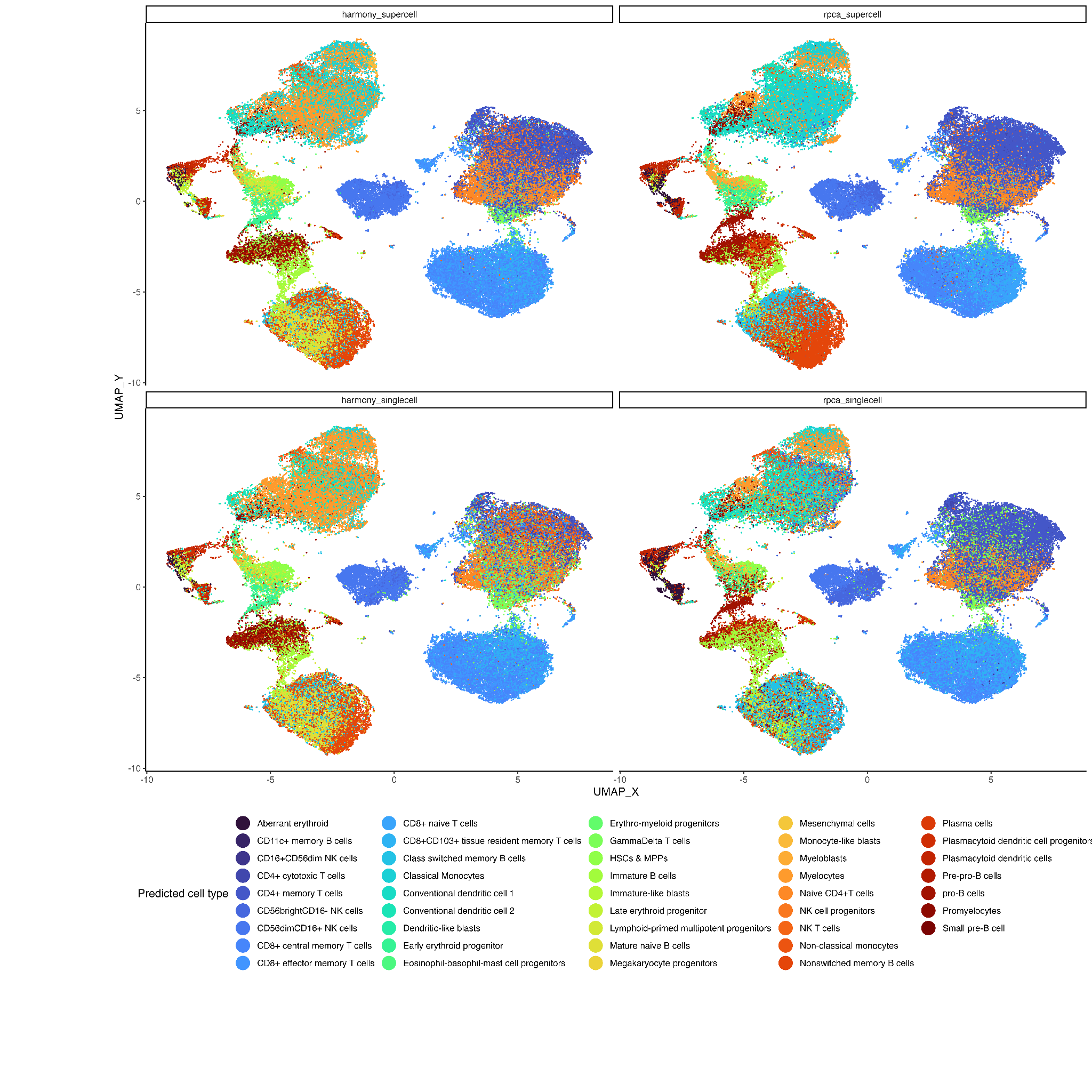

Supplementary Figure S17. UMAP plots of single cells for Levine_32dim dataset [[2]](https://www.zotero.org/google-docs/?jqdszk) annotated using either Harmony [[12]](https://www.zotero.org/google-docs/?wi7Jir) combined with a k-Nearest Neighbour (kNN) classifier or Seurat rPCA [[13]](https://www.zotero.org/google-docs/?OYQXyc) applied at either the single cell or the supercell level. Only cells assigned cell type labels using manual gating, i.e., the true label is known, are shown.

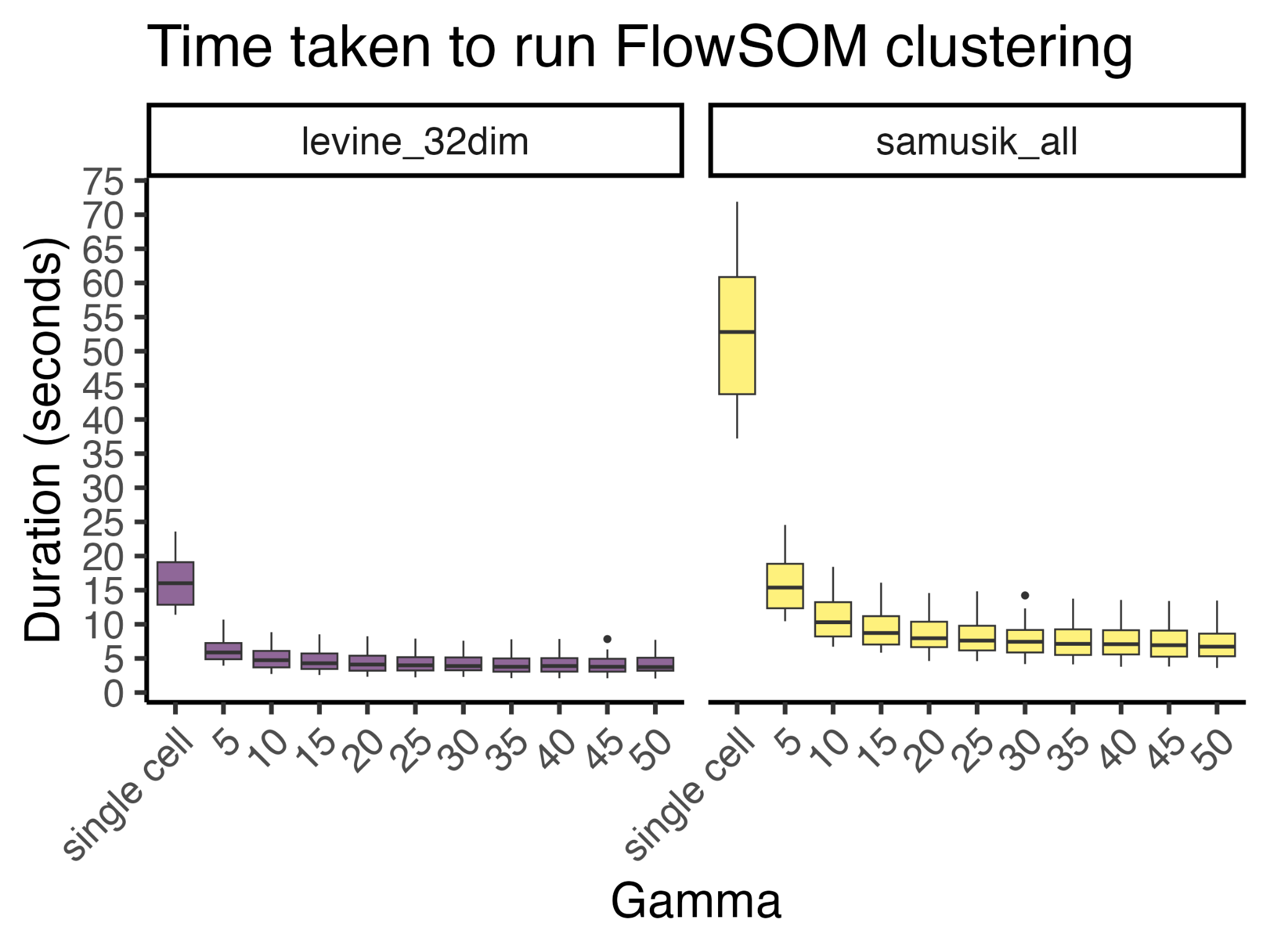

Supplementary Figure S18. The runtime of FlowSOM [[8]](https://www.zotero.org/google-docs/?WzXVmz) clustering for Levine_32dim and Samusik_all datasets, measured in seconds

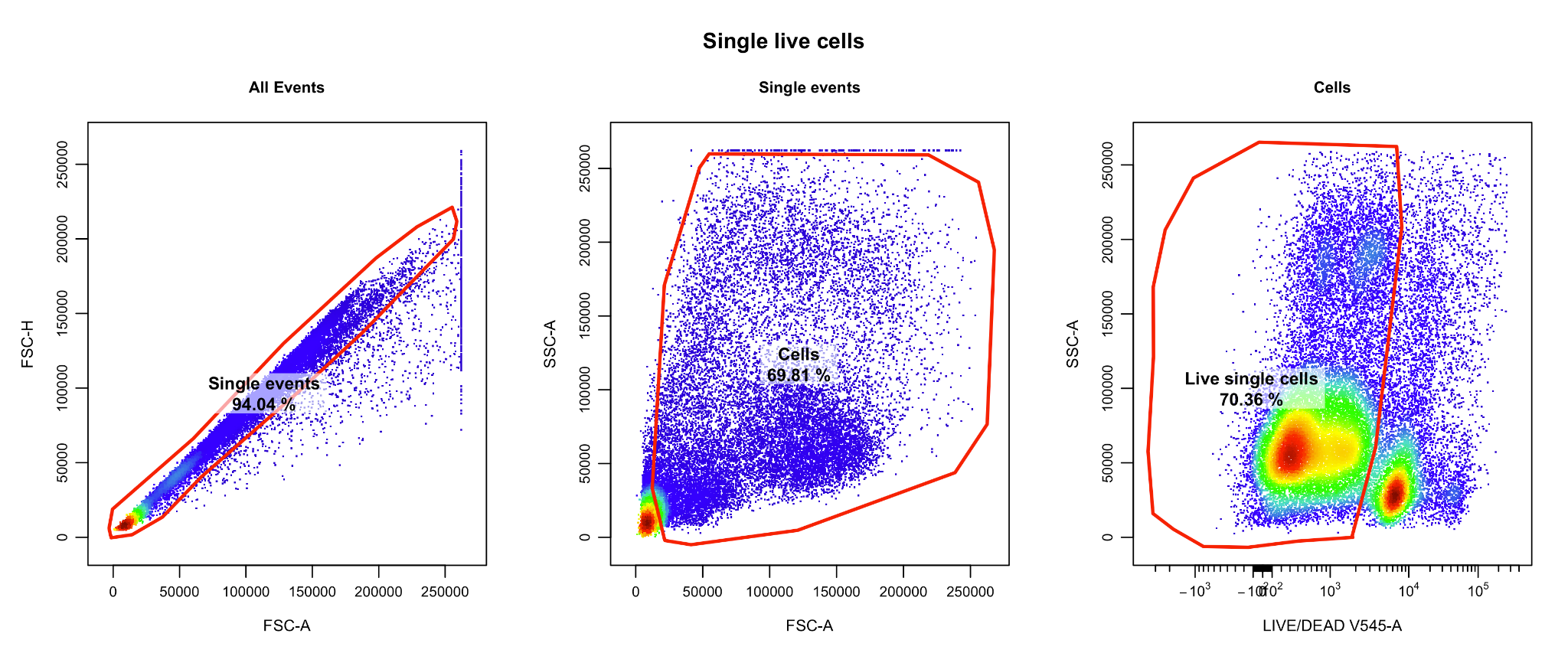

Supplementary Figure S19. The manual gating strategy used to isolate single live cells from the Oetjen_bcells [[1]](https://www.zotero.org/google-docs/?yCe4RE) flow cytometry dataset.
